# ER translocation of suboptimal targeting sequences depends on Sec61β/Sbh1 and its phosphorylation

**DOI:** 10.1101/2022.05.18.492448

**Authors:** Guido Barbieri, Julien Simon, Cristina R. Lupusella, Fabio Pereira, Francesco Elia, Hadar Meyer, Maya Schuldiner, Steven D. Hanes, Duy Nguyen, Volkhard Helms, Karin Römisch

## Abstract

The endoplasmic reticulum (ER) protein translocation channel subunit Sec61β/Sbh1 is non-essential, but contains multiple phosphorylation sites suggesting a regulatory role in ER protein import. We show here that mutating two N-terminal, proline-flanked, phosphorylation sites in the Sbh1 cytosolic domain phenocopies the temperature-sensitivity of a yeast strain lacking *SBH1/SBH2*, and results in reduced translocation into the ER of an Sbh1-dependent substrate, Gls1. In a microscopic screen we show that about 12% of GFP-tagged secretory proteins depend on Sbh1 for translocation. Sbh1-dependent proteins have targeting sequences with less pronounced hydrophobicity and often no or an inverse charge bias. A subset of these proteins was dependent on N-terminal phosphorylation of Sbh1 and on the phospho-S/T-specific proline isomerase Ess1 (PIN1 in mammals) for ER import. We conclude that Sbh1 promotes ER translocation of substrates with suboptimal targeting sequences and that its activity is regulated by a conformational change induced by N-terminal phosphorylation.

## INTRODUCTION

Protein secretion in eukaryotes starts with protein translocation across the endoplasmic reticulum (ER) membrane through the conserved Sec61 channel (*Mandon et al., 2013*; *O’Keefe and High, 2020*). The channel accommodates a myriad of different secretory and transmembrane proteins targeted to the ER via an N-terminal signal peptide (SP) or transmembrane domain (TMD) (*Mandon et al., 2013*; *O’Keefe and High, 2020*). Yeast also express an homolous channel, the Ssh1 channel, which has a distinct set of translocation substrates (*Mandon et al., 2013*). While many proteins are translocated into the ER constitutively, some are only required under specific circumstances. In addition, the concentration of ER chaperones and ER-resident enzymes has to be tightly controlled to maintain ER proteostasis (*Pilla et al., 2017*). This control is exerted at the transcriptional level by the Unfolded Protein Response (UPR), but for critical enzymes it may be prudent to also restrict or enhance their ER entry as an immediate response to altered physiological circumstances (*Pilla et al., 2017*; *Hegde and Kang, 2008*; *Hosokawa et al., 2010*).

ER SPs have a hydrophobic core of varying length, a net positive charge at the N-terminus that during translocation is oriented towards the cytosol, and a polar C-terminal region which contains the signal peptidase cleavage site (*O’Keefe and High, 2020*). They can have functions in addition to ER targeting including recruitment of cofactors necessary for their translocation through the Sec61 channel (*O’Keefe and High, 2020*). The Sec61 channel consists of three subunits (Sec61α, Sec61β, Sec61γ in mammals; Sec61, Sbh1, Sss1 in yeast), two of which (Sec61 and Sss1) are essential in yeast (*Mandon et al., 2013*). The 10 transmembrane helices of Sec61 form the channel itself with a hydrophobic constriction in the center (*Voorhees and Hegde, 2016*). During channel opening the entire N-terminal half of the channel including Sbh1 rotates by about 20 degrees to allow insertion of the translocation substrate into the lateral gate between TMDs 2 and 7 of Sec61 (*Voorhees and Hegde, 2016*). Sss1 consists of two helices which form a clamp around the Sec61 helix bundle that stabilizes the channel structure (*Voorhees and Hegde, 2016*).

Sec61β/Sbh1 is a tail-anchored protein that is peripherally associated with the channel via its conserved TMD that contacts TMDs 1 and 4 of Sec61 (*Mandon et al., 2013*; *Zhao and Jäntti, 2009*). Sec61β mediates interaction of signal peptidase and SRP receptor with the Sec61 channel (*Kalies et al., 1998*; *Jiang et al., 2008*). Sbh1 is central to the Sec complex required for posttranslational protein import into the yeast ER which consists of the Sec61 channel and the heterotetrameric Sec63 complex (Sec63, Sec62, Sec71, Sec72) (*Wu et al., 2019*). Although Sbh1 makes extensive contact with Sec71, it is dispensable for stability of the Sec complex and general posttranslational translocation into the yeast ER (*Allen et al., 2019*; *Bhadra et al., 2021*; *Feng et al., 2007*). The Sec61β/Sbh1 cytosolic domain consists of a membrane-proximal, conserved, and structured part of about 16 amino acids and an intrinsically unstructured, poorly conserved N-terminal domain of varying length (*Kinch et al., 2002*). It can bind ribosomes and the exocyst, but the role of these interactions is unclear (*Levy et al., 2001*; *Toikkanen et al., 2003*). In yeast, simultaneous deletion of *SBH1* and and the gene encoding its homolog in the Ssh1 channel, *SBH2,* results in temperature-sensitive growth (Fig. 1A), but the double deletion affects ER translocation of different substrates differentially (*Finke et al., 1996*; *Feng et al., 2007*). Although Sbh1 and Sbh2 are 53% identical at the amino acid level and both can interact with Sec61 or Ssh1, they also have distinct functions in translocation as shown by distinct patterns of synthetic lethality (*Toikkanen et al., 1996; Schuldiner et al., 2005; Jonikas et al., 2009; Costanzo et al., 2016*). Reconstitution of Sec61 channels lacking Sec61β into proteoliposomes still allows protein translocation, but only if the time for protein insertion is extended (*Kalies et al., 1998*). The Sec61β cytosolic domain makes contact with targeting sequences in the vestibule of the Sec61 channel, and this contact is enhanced if substrates are prevented from inserting into the lateral gate (*Laird and High, 1997*; *MacKinnon et al., 2014*). Taken together, the data suggest that Sec61β/Sbh1 recognizes some ER targeting sequences in the Sec61 channel vestibule and promotes their insertion into the lateral gate, but that its activity is not essential for most proteins.

**Figure 1.**
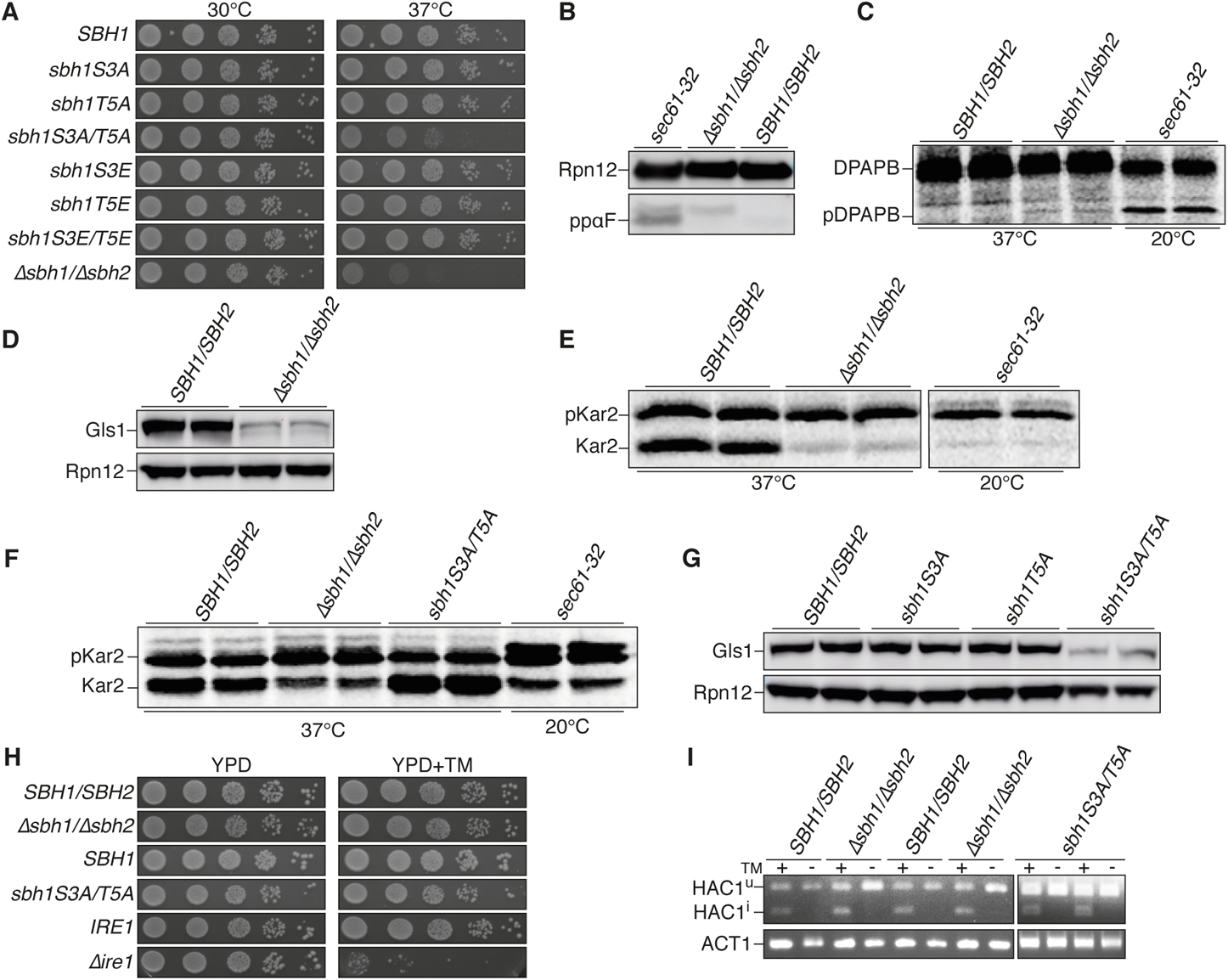
N-Terminal Sbh1 phosphorylation regulates transport of specific substrates into the ER. **(A)** Temperature-sensitivity test for *sbh1* mutants. The *Δsbh1Δsbh2* strain was transformed with a CEN-ARS vector expressing *SBH1* or the indicated *sbh1* mutants and serially diluted cells were grown at 30 °C or 37 °C for 3 days. **(B)** Wildtype, *Δsbh1Δsbh2,* and *sec61-32* strains grown to early exponential phase were lysed and post-translationally translocated ppαF or Rpn12 (loading control) were detected by Western blotting with specific antisera. **(C)** Wildtype and the indicated mutant strains were pulsed with [^35^S]-met/cys for 15 min and co-translationally translocated DPAPB was immunoprecipitated with specific antibodies. Cytosolic precursor (pDPAPB) and ER form (DPAPB) are indicated. **(D)** Cellular protein was extracted from wildtype and *Δsbh1Δsbh2* cells and Gls1 was detected by Western blotting with specific antibodies. Rpn12 was used as loading control. **(E; F)** Wildtype and the indicated mutant cells were pulsed with [^35^S]-met/cys for 2.5 min and Kar2 immunoprecipitated with specific antibodies. Cytosolic precursor (pKar2) and ER form (Kar2) are indicated. **(G)** Western blot analysis for Gls1 in *sbh1S3A/T5A* was done as in (D). **(H)** Wildtype (chromosomally *SBH1 SBH2*) and strains expressing the indicated genes from a CEN-ARS vector in *Δsbh1 Δsbh2* (as in A) were grown in serial dilutions on YPD and YPD+TM (0.5 μg/ml) plates for 3 days at 30 °C. The *Δire1* mutant was in a different background; the isogenic chromosomally *IRE1* wildtype is shown. **(I)** RNA was extracted from the indicated strains (as in H) grown without or with tunicamycin (TM) and an RT-PCR for *HAC1* and *ACT1* mRNA as control was performed followed by agarose gel-electrophoresis. Bands representing unspliced (HAC1^u^) and spliced (HAC1^i^) *HAC1* mRNA are indicated.

Transport of some proteins into the ER must be regulated, e.g. during specific developmental steps, under ER stress, or when pathogens encounter a host cell and need to secrete virulence factors. Regulation is possible if ER targeting sequences are not all equally strong, but if there are qualitative differences that determine the relative efficiency of insertion (*Hegde and Kang, 2008*; *O’Keefe and High, 2020*). In mammals many targeting sequences require accessory proteins to the Sec61 channel to accomplish translocation (*Hegde and Kang, 2008*; *O’Keefe and High, 2020*). Alternatively, function of the channel itself might be altered by posttranslational modifications (*Hegde and Kang, 2008*). A precedence is protein import into mitochondria where phosphorylation of both substrates and translocation machinery regulates import (*Opalinska and Meisinger, 2015*). The intrinsically unstructured domains of Sec61β and Sbh1 contain multiple phosphorylation sites, most of which are not positionally conserved (*Gruss et al., 1999*; *Soromani et al., 2012*). Mutation of all phosphorylation sites in Sbh1 individually to alanines (A), including the highly conserved, proline-flanked Threonine (T) in position 5, had no effect on the ability of the mutant *sbh1* to complement the temperature-sensitivity of a *Δsbh1Δsbh2* deletion strain (*Soromani et al., 2012*).

Progressive phosphorylation of intrinsically disordered domains, however, can act as a switch between one functional state and another, or induce formation of binding sites for specific interaction partners (*Valk et al., 2014*; *Bah and Forman-Kay, 2016*). We therefore asked whether we could identify groups of phosphorylation sites in Sbh1 that acted together on translocation into the ER, and targeting signals that were either generally Sbh1-dependent or dependent on Sbh1 phosphorylation. Mutating both N-terminal, proline-flanked phosphorylation sites in Sbh1 to A reproduced the temperature-sensitivity of a strain lacking both *SBH1* and *SBH2*. In a high content screen we identified about 12% of secretory proteins assayed as Sbh1-dependent. Having a broader list of affected proteins enabled us to uncover their commonalities. We found that Sbh1-dependent proteins had suboptimal ER targeting sequences, with lower hydrophobicity and frequently without or with an inverse charge bias. A small fraction of the screened proteins (2%) was dependent on N-terminal phosphorylation of Sbh1 and on the phospho-S/T-dependent proline isomerase Ess1 for translocation into the ER. We conclude that Sbh1 promotes ER import of substrates with suboptimal targeting sequences and that its activity can be regulated by a conformational change induced by N-terminal phosphorylation.

## RESULTS AND DISCUSSION

### N-Terminal Sbh1 phosphorylation regulates translocation of specific substrates into the ER

To investigate whether any combination of Sbh1 phosphorylation site mutations had an effect on *sbh1* function, we started with the two proline-flanked sites at Serine (S)3 and T5, (*Soromani et al., 2012*). We mutated S3 and T5 either to A or to Glutamic acid (E) (to mimic the phosphorylated site). All mutants promoted growth of the *Δsbh1Δsbh2* strain at the restrictive temperature, with the exception of the combination S3A/T5A, which resulted in reduced growth at 37 °C (Fig 1A). Our results suggest that the phosphorylation of both S3 and T5 together is important for Sbh1 function.

Previous data on the translocation defect in the *Δsbh1Δsbh2* strain were somewhat contradictory (*Finke et al., 1996*; *Feng et al., 2007*). To test whether the deletion of both *SBH1* and *SBH2* resulted in a general protein translocation defect, we investigated cytosolic precursor accumulation of ER translocation substrates. Post-translationally translocated pre pro alpha factor (ppαF) accumulated in a *sec61-32* mutant that has an ER import defect at 20 °C, but not in the *Δsbh1Δsbh2* strain at 37 °C (Fig 1B). Co-translationally translocated dipeptidylaminopeptidase B (DPAPB) also accumulated in the *sec61-32* mutant at the restrictive temperature, but not in the *Δsbh1Δsbh2* strain (Fig 1C). These results demonstrate that yeast cells do not have a general translocation defect either co-translationally or post-translationally in the absence of Sbh1 and Sbh2.

We previously reported a specific translocation defect for the cytosolic precursors of Gls1 and Mns1 (pGls1, pMns1) in the *Δsbh1Δsbh2* strain (Fig 1D; *Feng et al., 2007*). Finke *et al*. saw a moderate defect for Kar2 precursor (pKar2) translocation which was more pronounced in our hands (*Finke et al., 1996*) (Fig 1D, 1E, Fig EV1). We next investigated whether the *sbh1* S3A/T5A mutant was competent for translocation of these substrates. For testing pKar2 translocation, we performed a short pulse with [^35^S]-methionine (met)/cysteine (cys) and immunoprecipitation with Kar2-specific antibodies in wildtype, *Δsbh1Δsbh2*, *sbh1S3A/T5A,* and *sec61-32* mutant strains. We saw that in *sbh1S3A/T5A* yeast there was as much translocation of pKar2 as in the wildtype, in contrast to the *Δsbh1Δsbh2* and *sec61-32* strains, where we saw cytosolic pKar2 accumulation (Fig 1F, upper band), indicating that pKar2 is Sbh1-dependent, but not dependent on the S3/T5-phosphorylation of Sbh1. This was verified by Western blotting (Fig EV1). The cytosolic precursor (pGls1) and the ER form of Gls1 cannot be distinguished on SDS gels, but *Δsbh1Δsbh2* cells have a reduced amount of Gls1 in the ER at steady state (Fig 1D; *Feng et al., 2007*). We therefore quantified the amount of Gls1 in wildtype, *sbh1S3A/T5A,* and individual *sbh1S3A* and *sbh1T5A* strains by Western blotting. We found that the amount of Gls1 in the ER of *sbh1S3A/T5A* mutant was substantially reduced compared to the wildtype or the single mutants (Fig 1G), comparable to the reduction seen in *Δsbh1Δsbh2* strain (Fig 1D). This indicates that transport of Gls1 into the ER is dependent not only on the presence of Sbh1, but also on its phosphorylation at S3 and T5.

As Gls1, Mns1, and Kar2 are involved in ER protein quality control, we next investigated the sensitivity to the glycosylation inhibitor, Tunicamycin (TM), of *sbh1* mutants. Cells that are defective in ER-associated degradation (ERAD) or the unfolded protein response (UPR) like the *Δire1* mutant are sensitive to TM in the growth media (*Tran et al., 2011*) and TM is known to induce the UPR. We grew *Δsbh1Δsbh2*, *sbh1S3A/T5A,* and *Δire1* mutant strains, and the corresponding wildtypes on rich media either with or without TM (0.5 μg/ml). We found that in contrast to the *Δire1* strain, the *sbh1* mutants were not sensitive to TM (Fig 1H). We also tested these strains directly for induction of the UPR, by looking for the most proximal sign of induction - the splicing of the mRNA of the *HAC1* transcription factor mRNA - using as a positive control cells treated with TM. We did an RT-PCR for *HAC1* and *ACT1* mRNA as control. We found that neither the *Δsbh1Δsbh2* strain, nor the *sbh1S3A/T5A* mutant, in the absence of TM, contained spliced *HAC1* mRNA (Fig 1I), indicating that there is no induction of the UPR nor a proteostasis defect in the *sbh1* mutants. Our observations suggest that there are two classes of Sbh1-dependent ER translocation substrates: some are dependent on the presence of Sbh1, and a subset that are also dependent on S3/T5-phosphorylation of Sbh1.

Levels of glycan-processing enzymes like Gls1 and Mns1 and the molecular chaperone Kar2 in the ER need to be tightly controlled (*Hosokawa et al., 2003*). Our observations suggests that Sbh1 plays a role in the regulation of the ER import of these proteins under specific physiological circumstances. During active growth or induction of the UPR, ER import of these proteins would have to be maximized by phosphorylating Sbh1 (*Kabani et al., 2003; Hitt et al., 2004; Jakob et al., 1998*; *Simons et al., 1998*; *Orlean, 2012*), whereas in stationary phase or during recovery from the UPR their ER import needs to be limited by Sbh1 dephosphorylation to prevent excessive glycan-processing in the ER which would lead to disturbed ER proteostasis (*Hosokawa et al., 2003*).

### Identification of Sbh1-dependent and Sbh1-phosphorylation-dependent ER translocation substrates

To systematically identify proteins whose translocation depends on the presence of Sbh1 or the S3/T5-phosphorylation of Sbh1, we performed a high content screen (*Breker et al., 2014*). We integrated the *Δsbh1Δsbh2* or *sbh1S3A/T5A* backgrounds into a collection of yeast strains each expressing one of 382 secretory and transmembrane proteins C-terminally fused to a Green Fluorescent Protein (GFP) (*Geva et al., 2017*). We imaged the wildtype and mutant cells and analysed them for changes in the signal pattern. For example, we found that Pmt1, an ER-localized multispanning protein O-mannosyltransferase, exhibited an increase of the fluorescence signal in the *Δsbh1Δsbh2* mutant (Fig 2A, left vs. right). Another example is Msb2, an osmosensor involved in signal transduction with a single TMD that normally localizes to the vacuole, that displayed a reduction of the signal on the background of the *sbh1S3A/T5A* mutant (Fig 2B, left vs right), indicating a biogenesis defect. More globally, we identified 45 proteins that were dependent on the presence of Sbh1 and 7 proteins that where dependent on S3/T5-phosphorylation of Sbh1 (Table EV1); 5 of these were also identified in the Sbh1-dependence screen. To verify our results from the screens we biochemically analysed the translocation efficiency of two proteins that had an expected clear size difference between cytosolic precursor and ER form. Indeed, we found that for both Erp1-GFP and Gpi8-GFP there was cytosolic precursor accumulation in the *Δsbh1Δsbh2* strain, but not in wildtype or *sbh1S3A/T5A* mutant cells (Fig 2C).

**Figure 2.**
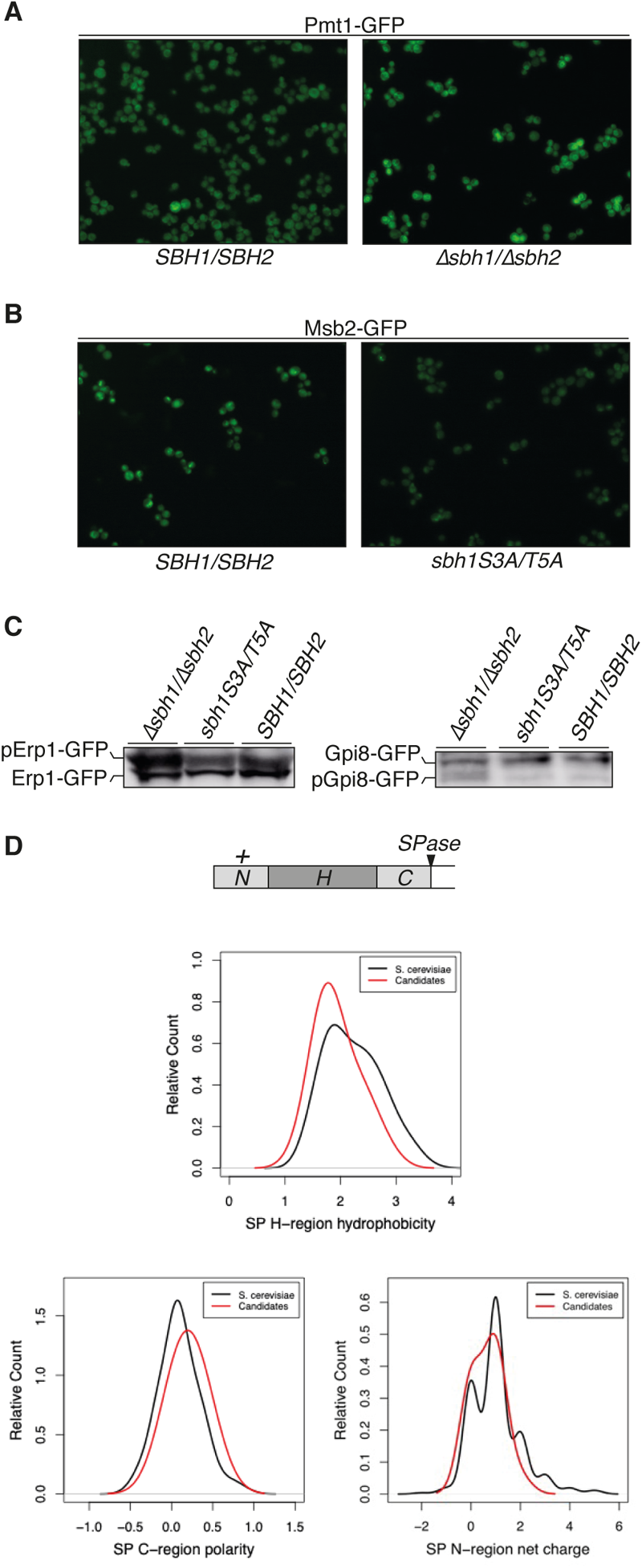
Identification of Sbh1-dependent and Sbh1 phosphorylation-dependent ER translocation substrates. **(A, C)** Image analysis from the high content screen of secretome-GFP library in wildtype cells and the indicated mutants. **(B)** Cellular protein was extracted from wildtype, and the indicated mutant cells and Erp1-GFP and Gpi8-GFP were detected by Western blotting with GFP antibodies. Cytosolic precursors (pErp1-GFP, pGpi8-GFP) and ER forms (Erp1-GFP, Gpi8-GFP) are indicated. **(D)** Schematic representation of a signal peptide, with a positively charged N-terminal region (N), hydrophobic region (H), C-terminal polar region (C), and signal peptide cleavage site (SPase). Physicochemical properties of signal peptides of Sbh1-dependent candidates (red) and total ER targeting sequences in yeast (black): average hydrophobicity of the core H-regions (top) according to the Kyte-Doolittle scale, ordered from less hydrophobic (left) to more hydrophobic (right), polarity of the C-regions (lower left), and net charge of N-regions (lower right).

SPs have a well-defined structure (Fig 2D, top). ER-targeting can also be achieved by uncleaved SPs (signal anchors, SAs) or the first TMD of a protein (*Spiess et al., 2019*). Identification of a significant number of Sbh1-dependent proteins allowed us to investigate whether their ER targeting sequences had specific common features compared to the total ER targeting sequences in yeast. Several features make a SP optimal for ER translocation through the Sec61 channel: Helix propensity, which can be disturbed by high glycine/proline content (*Nguyen et al., 2018*); hydrophobicity of the H-region, with a great diversity in terms of length; and charge bias between N-region and C-region, which helps the peptide orientate when inserting to the channel are among the most important ones (*Spiess et al., 2019*; *Yim et al., 2018*). Of the 45 Sbh1-dependent proteins, 16 had SPs, 5 had SAs, and 24 had TMDs as ER targeting signals (Table EV1). We found that the Sbh1-dependent SPs were slightly less hydrophobic (Fig 2D, top graph), but we detected no differences in polarity of the C-region or charge distribution (Fig 2D, lower panels). When we looked at Sbh1-dependent ER targeting sequences individually, however, we found that many targeting sequences had no charge bias (e.g., Yps7, Fig EV2A) or an inverse charge bias (e.g., Gpi14, Fig EV2A). This was true for both SPs and transmembrane targeting sequences of Sbh1-dependent proteins. In addition, some transmembrane targeting sequences were unusually long or too short to span the membrane (e.g., Yip3, Fig EV2B) or contained a high number of glycine residues (e.g., Tat1, Fig EV2B); all of these features would interfere with the efficient insertion of these targeting sequences into the lateral gate of the Sec61 channel (*Nguyen et al., 2018*; *Spiess et al., 2019*; *Yim et al., 2018*). Targeting sequences of Sbh1 S3/T5-phosphorylation-dependent proteins were similar to the Sbh1-dependent ones (Fig EV2C), but we were unable to identify specific features in these due to the small number of proteins identified. Our observations suggest that Sbh1-dependent proteins have ER-targeting sequences that are suboptimal for insertion into the Sec61 lateral gate. Sbh1 may guide these targeting sequences into the Sec61 channel and thus enhance their insertion efficiency.

### Sbh1 is required for cell wall biogenesis

Since many of the Sbh1-dependent proteins that we found play a role in cell wall biogenesis (Table EV1, blue), we investigated whether the *Δsbh1Δ*sbh2 mutants had a cell wall defect. We grew *Δsbh1Δsbh2* and the *sec61-3* mutant as a positive control alongside the corresponding wildtype strains on rich media (YPD) and YPD supplemented with 1.2M sorbitol (YPDS), which stabilizes the plasma membrane if the cell wall is defective (*Lommel et al., 2004*), at 30 °C and 37 °C. We found that both the *Δsbh1Δsbh2* strain and the *sec61-3* mutant grew at 37 °C in the presence of sorbitol (Fig 3A), suggesting that it is indeed the cell wall defect that makes the Sbh1/2 mutant cells temperature sensitive. In addition, we found that Sbh1 expression and Sbh1 phosphorylation is considerably higher in early exponential phase, compared to later stages (Fig EV3), consistent with its requirement for cell wall biosynthesis.

**Figure 3.**
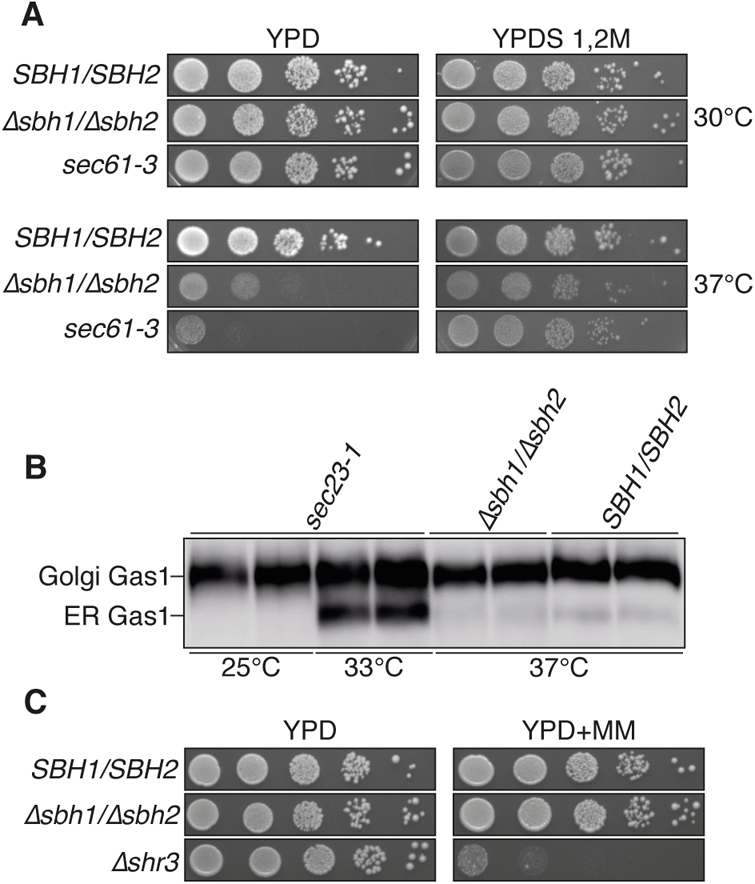
Sbh1 is required for cell wall biogenesis. **(A)** Wildtype and the indicated mutant strains were grown in serial dilutions on YPD and YPD+1.2 M sorbitol (YPDS) for 3 days at 30 °C and 37 °C. **(B)** Cellular protein was extracted from wildtype and the indicated mutant strains and Gas1 was detected by Western blotting with specific antibodies. *Sec23-1* cells were grown for 3 h at the restrictive temperature (33 °C) prior to lysis. Wildtype and *Δsbh1Δ*sbh2 cells were grown for 3 h at 37 °C prior to lysis. Golgi and ER Gas1 forms are indicated. **(C)** Wildtype and the indicated mutant strains were grown in serial dilutions on YPD and YPD+metsulfuron-methyl 200 μg/ml (YPD+MM) plates for 3 days at 30 °C.

As several of the Sbh1-dependent proteins are involved in the biosynthesis of glycosylphosphatidylinositol anchors (GPI-anchor) that are glycolipid anchors enabling the presentation of proteins on the outer membrane or cell wall (Table EV1, blue), we asked whether the *Δsbh1Δsbh2* strain was affected in GPI-anchor synthesis. The GPI-anchored protein Gas1 accumulates in the ER if its GPI-anchor is not properly processed (*Horvath et al., 1994*). Cells lacking *SBH1* and *SBH2*, however, did not accumulate the ER form of Gas1 after a 3 h shift to the restrictive temperature (Fig 3B). Our data suggest that despite the reduction of components of the GPI-synthesis machinery in the ER of *Δsbh1Δsbh2* mutants, the strain was competent for GPI anchor production.

Since several of the Sbh1-dependent proteins found in our screen are amino acid transporters in the plasma membrane (Table EV1, red), we investigated whether the *Δsbh1Δ*sbh2 mutant had a defect in amino acid uptake. We tested its ability to survive on plates supplemented with metsulfuron-methyl (MM), which is toxic in strains lacking amino acid transporters (*Jørgensen et al., 1998*). In contrast to an *Δshr3 mutant* (chaperone, required for amino acid transporter biogenesis, *Kuehn et al., 1998*), we found that *Δsbh1Δsbh2* was not sensitive to MM (Fig 3C), suggesting that although amino acid transporter biogenesis was reduced in these cells, amino acid uptake was not critically affected.

Both *Δsbh1Δsbh2* and the *sec61-3* strain have a cell wall defect that results in temperature-sensitivity. Since a large fraction of *S. cerevisiae* secretory activity is dedicated to cell wall biogenesis (*Guo et al., 2021*) this makes it likely that the temperature-sensitivity of most if not all secretory pathway mutants is due to an underlying cell wall defect (*Novick and Schekman, 1983*).

### Screening for the kinase responsible for Sbh1 S3/T5 phosphorylation

To identify the kinase responsible for the phosphorylation of proline-flanked S3/T5 of Sbh1, we raised an antibody against the phosphorylated N-terminus of Sbh1 (Sbh1_(Pi)_), that recognizes N-terminally phosphorylated Sbh1 better than the unphosphorylated form (Fig 4A, right panel). Initially, we used the Sbh1_(Pi)_ antibody vs. the Sbh1_(10-23)_ antibody that recognizes N-terminally unphosphorylated Sbh1 better than its phosphorylated counterpart (Fig 4A, left) to screen through loss of function mutants in all 27 proline-directed kinases in yeast (Table EV2, *Kanshin et al., 2017*; *Zhang et al., 2019*; *Zhu et al., 2005*; *Bradley and Beltrao, 2019*; *Liu et al., 2000*; *Lin et al., 1997*). We were unable to identify a kinase mutant in which N-terminal Sbh1 phosphorylation was reduced. We also used the antibodies to screen for Sbh1 N-terminal hyperphosphorylation (*Sopko et al., 2006*) in strains overexpressing 20 of these proline-directed kinases, again without a conclusive result.

**Figure 4.**
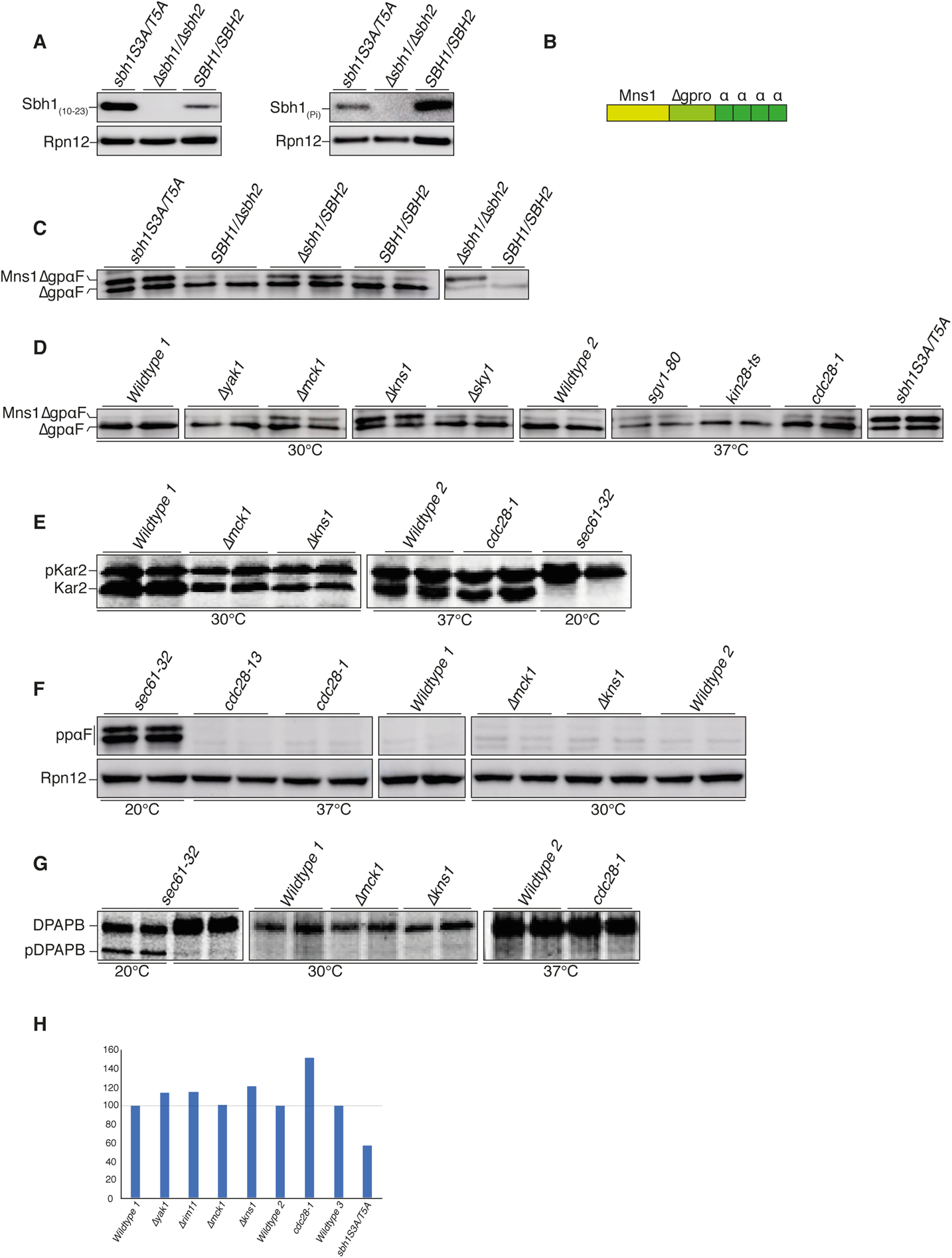
Screening for the kinase responsible for Sbh1 S3/T5-phosphorylation. **(A)** Wildtype, *Δsbh1Δsbh2,* and *sbh1S3A/T5A* strains grown to early exponential phase were lysed and Sbh1 or phosphor (Pi)-Sbh1 were detected by Western blotting with antibodies against amino acids 10-23 of Sbh1 (Sbh1_(10-23)_, left) or the phosphorylated N-terminus (Sbh1_(Pi)_, right). Note that the Sbh1_(10-23)_ antibody preferentially recognizes N-terminally unphosphorylated Sbh1, whereas the Sbh1_(Pi)_ antibody preferentially recognizes N-terminally phosphorylated Sbh1. Rpn12 was used as loading control. **(B)** Schematic representation of the Sbh1 S3/T5 phosphorylation-dependent reporter construct Mns1ΔgpαF. Mns1 signal peptide (yellow), mutant pro region lacking N-glycosylation sites (light green), alpha factor repeats (α)are indicated. **(C, D, E)** Cellular protein was extracted from wildtype and the indicated mutant cells grown to early exponential phase and Mns1ΔgpαF was detected by Western blotting with specific antisera. **(F)** Wildtype and the indicated mutant cells were pulsed with [^35^S]-met/cys for 2.5 min and Kar2 immunoprecipitated with specific antibodies. Cytosolic precursor (pKar2) and ER form (Kar2) are indicated. **(G)** Cellular protein was extracted from wildtype and the indicated mutant cells and post-translationally translocated ppαF was detected by Western blotting with specific antisera. **(H)** Wildtype and the indicated mutant strains were pulsed with [^35^S]-met/cys for 15 min and co-translationally translocated DPAPB was immunoprecipitated with specific antibodies. Cytosolic precursor (pDPAPB) and ER form (DPAPB) are indicated. **(I)** Cellular protein was extracted from wildtype and indicated mutant cells and Gls1 was detected by Western blotting with specific antibodies and quantified in duplicates relative to the loading control Rpn12.

We then generated a reporter construct, fusing the SP of the Sbh1 phosphorylation-dependent substrate Mns1 to mutant alpha factor precursor without glycosylation sites (Mns1ΔgpαF, Fig 4B). We first characterized the construct in our *sbh1* mutants. We detected reporter precursor accumulation in *Δsbh1Δsbh2*, *Δsbh1/SBH2,* and *sbh1S3A/T5A* strains (Fig 4C, upper band). We transformed the 27 proline-directed kinase deficient mutants (knockout non-essential, temperature sensitive for essential) and their respective wildtypes with a vector expressing this reporter construct (Mns1ΔgpαF), and found some cytosolic precursor accumulation in *Δkns1*, *Δmck1*, and in temperature-sensitive *cdc28-1* at the restrictive temperature (Fig 4D; Fig EV4A). These kinases might therefore be involved in Sbh1 N-terminal phosphorylation.

We subsequently investigated translocation of the Sbh1-dependent, but N-terminal phosphorylation-independent, translocation substrate pKar2 in these kinase mutants in pulse-assays. None of the kinase mutants had defects in pKar2 translocation (Fig 4 E), suggesting that the observed effect with the reporter was specific to the Mns1 SP. Both post-translational import of wildtype ppαF (Fig 4F) and co-translational import of pDPAPB (Fig 4G) was functional in these kinase mutants and we did not see any precursor accumulation, suggesting the kinase mutations do not cause general translocation defects.

We then investigated whether the proline-directed kinase mutants affected the levels of endogenous Gls1 in the ER. In contrast to the sbh1 S3A/T5A mutation which reduces Gls1 to around 50% of wildtype levels (Fig 4H), none of the kinase mutants identified in our screen significantly affected Gls1 levels in the ER of the mutants (Fig 4H). In addition, we investigated Gls1 translocation in these cells with a reporter that had glutathione-S-transferase fused to the N-terminus of the Gls1 SP (Fig EV4C) to generate a protein with a more pronounced size difference between cytosolic precursor and signal-cleaved ER-form (*Shibuya et al., 2015*). While *sbh1S3A/T5A* cells accumulated significant amounts of GSTpGls1 in the cytosol (Fig EV4D), none of the proline-directed kinase mutants did (Fig EV4E). Our results suggest that despite their defect in Mns1ΔgpαF translocation *Δkns1*, *Δmck1*, and *cdc28-1* cells were competent for Gls1 import into the ER and hence none of these kinases is likely responsible for N-terminal Sbh1 phosphorylation.

There are a number of possible reasons for our inability to identify the Sbh1 S3/T5 kinase: one possibility is the adaptation of the mutant strain to the absence of the kinase. Adaptation has been observed in yeast strains deficient in SRP function (*Ogg et al., 1992*). Also, kinase redundancy is common in yeast and has been reported e.g., for Fus3 and Kss1 targets (*Elion et al., 1991*).

### Phosphorylation-dependent proline isomerase Ess1 contributes to Sbh1 regulation

Both S3 and T5 in Sbh1 are proline-flanked (Fig 5A). The conserved enzyme Ess1 (PIN1 in mammals) isomerizes proline residues that are preceded by phosphorylated serine or threonine (Fig 5B), so the phosphorylated N-terminus of Sbh1 is a potential Ess1 target (Hanes et al., 2014). An active site mutant, *ess1H164R*, is synthetically lethal with *ssh1* indicating a contribution of Ess1 to ER protein translocation (*Gemmill et al., 2005*; *Atencio et al., 2014*). In membrane fraction experiments we found that about 30% of both wildtype and mutant *Ess1* was associated with a crude yeast microsome fraction, suggesting that Ess1 has membrane-bound targets (Fig 5C).

**Figure 5.**
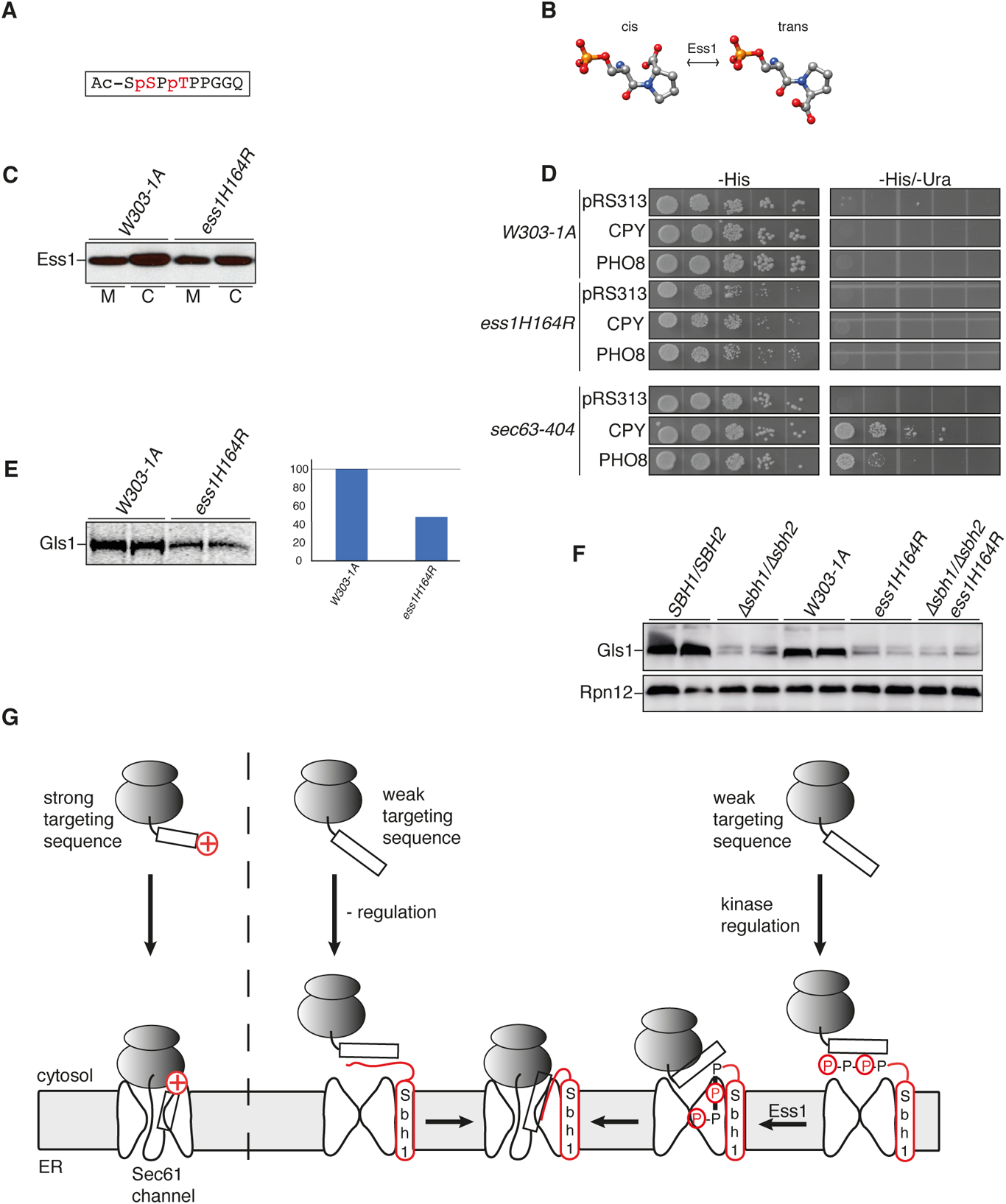
Phosphorylation-dependent proline isomerase Ess1 contributes to Sbh1 regulation. **(A)** Schematic representation of the phosphorylated-S3/T5 Sbh1 N-terminus. Phosphorylated amino acids shown in red. **(B)** Ess1-catalyzed phosphoserine-proline isomerization. Carbon atoms in grey, oxygen atoms in red, nitrogen atoms in blue, phosphorous atom in orange, hydrogen atoms not shown. **(C)** Wildtype and *ess1H164R* strains were grown to early exponential phase, lysed, and membranes sedimented; proteins were resolved by SDS-PAGE and Ess1 was detected by Western blotting with specific antisera. **(D)** Wildtype and the indicated mutant strains were transformed with a vector expressing ER translocation reporter constructs, with the signal peptide of post-translationally translocated CPY or co-translationally translocated Pho8 fused to the *URA3* gene and grown in serial dilutions on - His and -His/-Ura plates for 3 days at 30 °C. As control strains transformed with the empty vector (pRS313) were used. **(E)** Wildtype and the indicated mutant cells were pulsed with [^35^S]-met/cys for 5 min and Gls1 was immunoprecipitated with specific antibodies. Quantitation of Gls1 on the right. **(F)**. **(G)** Cellular protein was extracted from wildtype and the indicated mutant strains and Gls1 was detected by Western blotting with specific antibodies. Rpn12 was used as loading control. **(H)** Model for Sbh1 function during ER protein translocation and its regulation by S3/T5 phosphorylation.

To investigate whether the *ess1H164R* mutant had any ER translocation defects we used reporter constructs in which the SP of post-translationally translocated CPY or co-translationally translocated Pho8 were fused to the *URA3* gene (*Ng et al., 2007*). When these are expressed in *ura3* auxotrophs, the cells can only survive in the absence of uracil if the reporter fails to translocate into the ER or does so more slowly (*Ng et al., 2007*). Using this assay, we found that *ess1H164R* does not cause general translocation defects (Fig 5D, upper panels), in contrast to our control, *sec63-404*, which has a strong post-translational and a weaker, but still detectable, co-translational translocation defect (*Servas and Römisch, 2013*) (Fig 5D, lower panels). The *ess1H164R* mutant also had no effects on pKar2 import into the ER in a pulse experiment (Fig EV2A).

We next investigated whether the *ess1H164R* mutant was competent for translocation of the S3/T5 Sbh1-phosphorylation-dependent Gls1. We performed a pulse with [^35^S]-met/cys for 5 min and immunoprecipitated Gls1 with specific antibodies from wildtype and *ess1H164R* mutant cell lysates. We found that translocation of Gls1 into the ER of *ess1H164R* cells was reduced to about 50% of the wildtype (Fig 5E), comparable to the reduction seen in the *sbh1S3A/T5A* strain (Fig 1G). This indicates that transport of Gls1 into the ER is dependent not only on the phosphorylation at S3 and T5 of Sbh1, but also on the isomerization by Ess1. In addition, we found that temperature-sensitivity of the *ess1H164R* mutant at 35 °C was suppressed in the presence of sorbitol (Fig EV5B) confirming prior results by Gemmill *et al*. (2005) and suggesting that a cell wall defect, similar to the one found in the *Δsbh1Δsbh2* mutant strain (Fig 3A), makes the *ess1H164R* mutant temperature-sensitive.

To investigate whether Ess1 and Sbh1 S3/T5 phosphorylation control the same step in ER translocation, we tested whether the effect on Gls1 translocation in *sbh1S3A/T5A* and *ess1H164R* was additive. We measured the amount of Gls1 in *Δsbh1Δsbh2, ess1H164R,* a triple mutant containing *Δsbh1Δsbh2* and *ess1H164R,* and their respective wildtype strains by Western blotting. We found that the amount of Gls1 in the ER of all mutant cells was similarly reduced compared to wildtype (Fig 5F), suggesting that Ess1 and Sbh1 operate at the same stage of translocation.

Our data suggests the following model for ER protein translocation regulation by Sbh1 N-terminal phosphorylation and Ess1: When a ribosome-nascent chain complex with a suboptimal targeting sequence arrives at the Sec61 channel, failure to insert into the lateral gate leads to contact of the targeting sequence with the Sbh1 cytosolic domain in the Sec61 channel vestibule (Fig 5G, centre). Interaction with Sbh1 allows the targeting sequence to acquire the appropriate conformation, orientation or both for insertion into the lateral gate (Fig 5G, centre). For proteins whose concentration in the ER needs to be tightly controlled, phosphorylation of S3/T5 and isomerization by Ess1 enhance Sbh1-promoted ER import under specific physiological circumstances (Fig 5G, right). During active growth, e.g., ER import of Mns1 and Gls1 precursors would be maximal, whereas in stationary phase or during recovery from the UPR their ER import might be limited to prevent excessive glycan-processing in the ER which would lead to disturbed ER proteostasis. When cells are exposed to an increased environmental osmolarity, ER import would be maximal for osmosensors and for proteins that are involved in the high osmolarity response (HOG) pathway, contributing to hyperosmotic stress tolerance, like Vph1 and Msb2 (two other phospho-Sbh1 dependent substrates (Table EV1)). In addition, the HOG pathway plays a collaborative role with the Cell Wall Integrity (CWI) pathway, by inducing cell wall remodelling (*Udom et al., 2019*).

As shown for other intrinsically disordered regions, phosphorylation of the N-terminus of Sbh1 may affect protein conformational dynamics or liquid-liquid phase separation (*Bah and Forman-Kay, 2016*), thus regulating interaction with specific ER targeting sequences. Our results indicate that access to the ER of these substrates is further controlled by Ess1-dependent isomerization. More generally our results demonstrate how intricate the ER protein translocation system is, enabling the tight regulation and tailoring of translocation according to cellular needs.

## MATERIALS AND METHODS

### S. cerevisiae strains

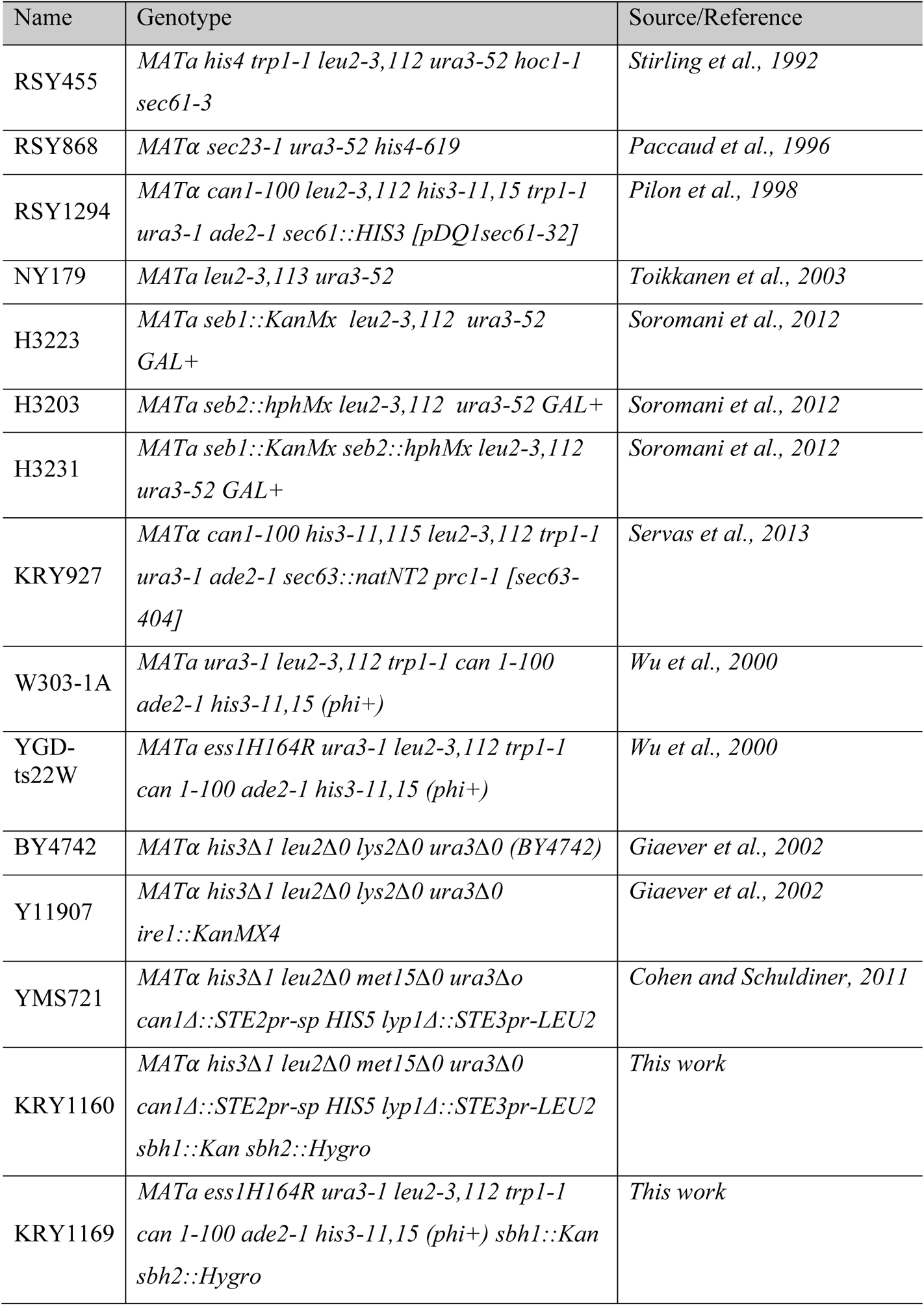
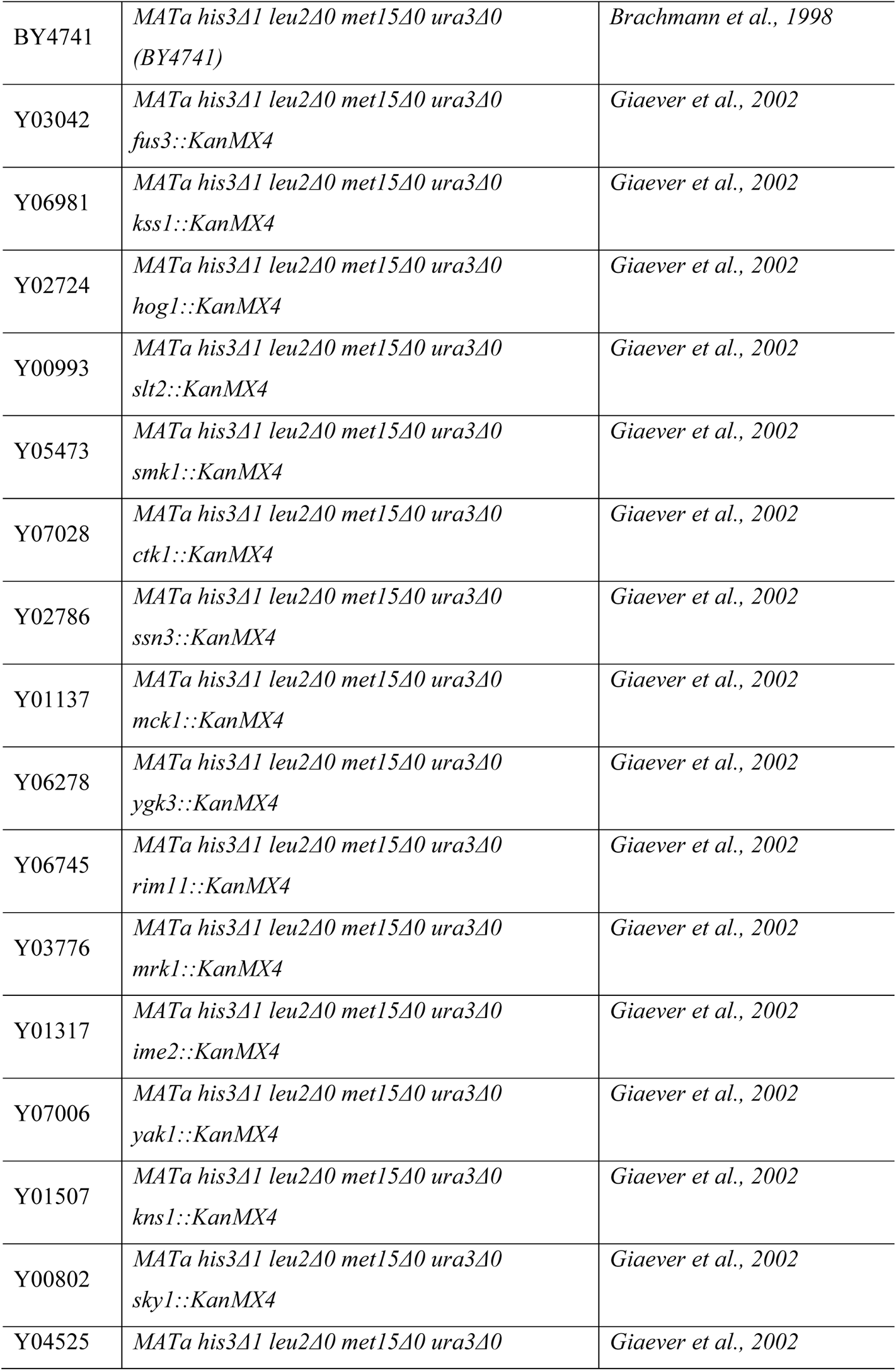
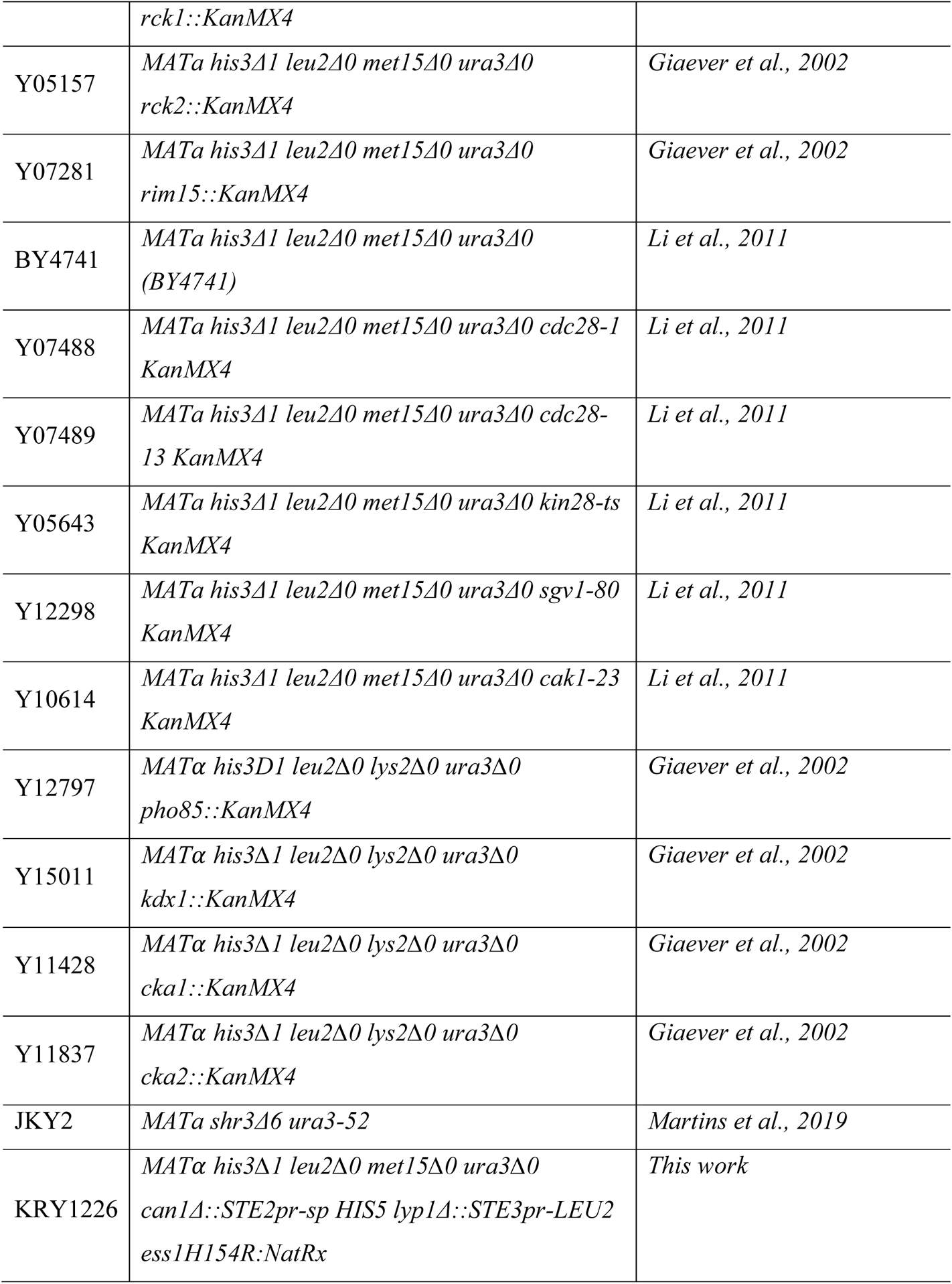

Please note that *SBH1* was originally named *SEB1*, and *SBH2* was *SEB2* (Toikaanen et al., 1996).

The library used for the screens was the mini “Secretome-GFP” library, genotype: BY4741 *Δmet Δura Δhis Δleu MATa XXX-GFP-HIS* (*Huh et al., 2003*).

The strains overexpressing 20 of the proline-directed kinases to screen for Sbh1 N-terminal hyperphosphorylation are form the TEF2-Cherrry Overexpression library, genotype: Δura Δhis Δleu Δmet MATa Δcan1::STE2pr-spHIS5 Δlyp1::STE3pr-LEU2 NATR-TEF2pr-mCherry-XXX (Weil et al., 2018)

### Antibodies

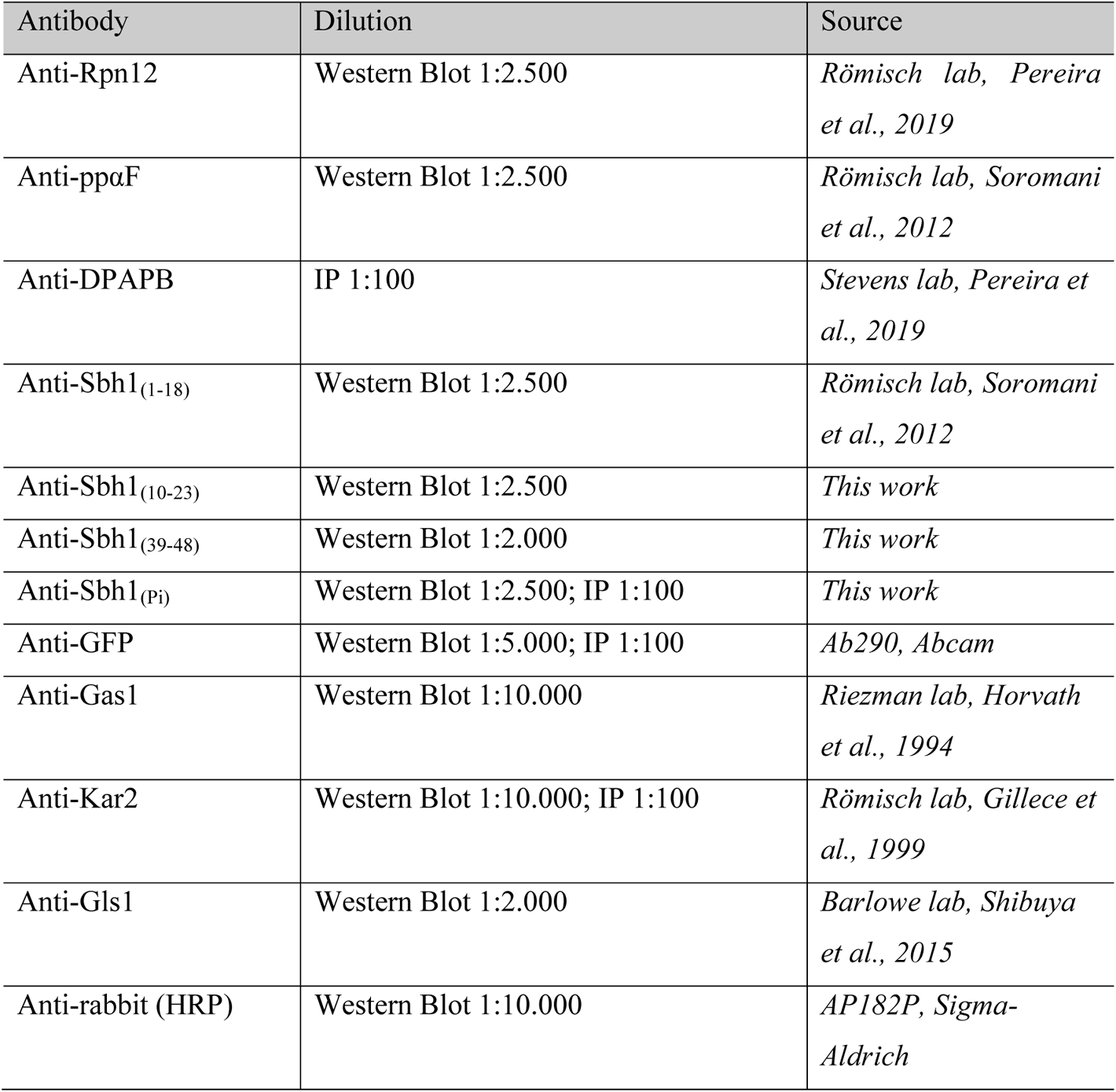

### Growth of S. cerevisiae

Yeast strains were grown either in full or selective media at 30 °C (if not stated otherwise) with continuous shaking at 220 rpm, and cells were harvest in early exponential phase and washed with sterile deionized water. For drop dilution assays an OD_600_ of 0.5 was harvested, washed and serial 1:10 dilution was done. For each dilution, 5 μl (containing 10^4^-10 cells) were spotted on to the respective media plates. To test tunicamycin (TM) (Sigma) sensitivity, cells were grown on YPD plates supplemented with 0.5 μg/ml TM. To test metsulfuron-methyl (MM) (Sigma) sensitivity, cells were grown on YPD plates supplemented with 200 μg/ml MM. To test the effect of sorbitol on growth recovery, cells were grown on YPD plates supplemented with 1.2 M sorbitol. The growth was documented after 3 days.

### Protein Extraction

Yeast strains were grown to an OD_600_ of 1 at 30 °C, 220 rpm. Cells 2 (OD_600_) were harvested at 1.600 x g for 1 min and the supernatants were discarded. Pellets were washed with 1 ml of sterile deionized water, resuspended in 200 μl 2X SDS sample buffer (100 mM Tris-HCl, pH 6.8/4% SDS/0.2% bromophenol blue/20% glycerol/200 mM DTT). Glass beads (100 μl, acid washed, 1 mm, Sigma) were added and the cells were disrupted in a MiniBeadbeater-24 (Bio Spec Products Inc.) at 4 °C for 2 x 1 min, with 1 min pause in between cycles. Samples were incubated at 95 °C for 5 min (10 min at 65 °C for membrane proteins) and centrifuged at 11.000 x g for 1 min. Samples were then loaded onto SDS-PAGE gels.

### Isolation of Membrane sand Cytosolic Fractions

Yeast strains were grown to early exponential phase at 30 °C, 220 rpm. Cells (7 OD_600_) were harvested at 2.000 x g for 5 min and washed with 1 ml of 100 mM Tris-HCl, pH 9.4. After addition of 10 mM DTT, cells were incubated at RT for 10 min, and centrifuged at 4.300 x g for 1 min. Pellets were resuspended in 200 μl of JR lysis buffer (20 mM HEPES, pH 7.4/50 mM KOAc/2 mM EDTA, pH 8/1 mM DTT/1 mM PMSF/1x Phosphatase inhibitor cocktail, Thermo Scientific), 0.3 g of glass beads (acid washed, 1 mm, Sigma) were added and the cells were disrupted in a MiniBeadbeater-24 (Bio Spec Products Inc.) at 4 °C for 2 x 1 min, with 1 min pause in between cycles. After short centrifugation (10 sec, 14.000 x g), the supernatant was transferred to a to a clean microfuge tube and the glass beads were washed with 100 μl B88 (20 mM HEPES, pH 6.8/250 mM sorbitol/150 mM KOAc/5 mM Mg(OAc)_2_/1x Phosphatase inhibitor cocktail, Thermo Scientific) and the new supernatant was added to the previous one. Membranes were sedimented for 15 min, 14.000 x g, 4 °C. The supernatant corresponds to the cytosolic fraction and was transferred to a clean microfuge tube. The final volume was adjusted to 350 μl with 2X SDS sample buffer (100 mM Tris-HCl, pH 6.8/4% SDS/0.2% bromophenol blue/20% glycerol/200 mM DTT). Sedimented membranes were used for alkaline phosphatase treatment and subsequent TCA precipitation.

### Alkaline Phosphatase Treatment and TCA Precipitation

Sedimented membranes were resuspended in 50 μl B88 (20 mM HEPES, pH 6.8/250 mM sorbitol/150 mM KOAc/5 mM Mg(OAc)_2_/1x Phosphatase inhibitor cocktail, Thermo Scientific) and 20 μl of Alkaline Phosphatase (1u/μl, FastAP, Thermo Scientific) was added together with 8 μl of the reaction buffer (10x Thermo scientific FastAP reaction buffer). The samples were incubated for 1h at 37 °C. Membranes were then sedimented at 20.000 x g at 4 °C for 10 min, and resuspended in 100 μl B88. Membranes were sedimented as before and resuspended in 100 μl B88. Samples were then ready for TCA precipitation. As non-AP treated control, sedimented membranes were directly resuspended in 100 μl B88, without any AP treatment. Proteins were precipitated with 20% TCA on ice for 30 min and washed with ice-cold acetone. After centrifugation of the samples for 5 min, 14.000 x g, 4 °C, pellet was resuspended in 140 μl 2x SDS sample buffer (100 mM Tris-HCl, pH 6.8/4% SDS/0.2% bromophenol blue/20% glycerol/200 mM DTT) and incubated at 65 °C for 10 min.

### Immunoblotting

Protein gel electrophoresis was conducted using Bolt pre-cast Bis-Tris Plus gels (4-12%, 1 mm). Proteins were transferred to nitrocellulose membranes (BioRad) and detected with specific antibodies at the appropriate dilutions, and an ECL reagent (Thermo Scientific) according to the supplier’s instructions. Signal was acquired using an Amersham Imager 600 (GE Healthcare).

### Isolation of RNA

Yeast strains (10 ml culture) were grown to early exponential phase at 30 °C, 220 rpm. Cells were harvested for 5 min at 4 °C, 5.700 x g. Pellets were resuspended in 1 ml of ice-cold RNase-free water (DEPC-treated). The cells were centrifuged at full speed, 4 °C for 10 sec (5424-R Eppendorf microfuge) and the pellet was resuspended with 400 μl TES Solution (10 mM Tris-HCl, pH 7.7/10 mM EDTA/0.5% SDS). Acid Phenol (400 μl, Roti®-Aqua-Phenol, Carl Roth) was added and the sample vortexed for 10 sec and incubated at 65 °C for 1 h with occasional vortexing. The samples were centrifuged at full speed, 4 °C for 5 min. The aqueous phase was transferred to a clean microfuge tube. Roti®-Aqua-Phenol (400 μl) was added and the sample was vortexed for 20 sec, incubated on ice for 5 min and centrifuged as before. The aqueous phase was transferred to a clean microfuge tube and mixed with 400 μl chloroform, vortexed for 20 sec and centrifuged as before. The aqueous phase was transferred to a clean microfuge tube and 40 μl of 3 M sodium acetate (pH 5.3) and 1ml of ice-cold 100% ethanol were added. The sample was vortexed and centrifuged as before. Pellets were washed with 1.5 ml 70% ethanol, centrifuged as before and resuspended in 50 μl RNase-free water (DEPC-treated).

### HAC1 mRNA Splice Assay

Yeast strains were grown to early exponential phase at 30 °C, 220 rpm. For positive controls each strain was incubated in the presence of TM (2 μg/ml), 3 h as above. A volume of 10 ml of each culture was pelleted, and used to isolate yeast RNA. The RNA was then diluted to a final concentration of 0.1 μg/μl, and used in reverse transcription reactions to generate cDNA using the Maxima® Reverse Transcriptase (Fermentas), according to the manufacturer’s instructions. Each cDNA (0.1 μg) was used in a PCR reaction using *HAC1* and *ACT1* specific primer sequences. The PCR products were resolved on a 1% agarose gel. Bands were visualized and photographed using the E-BOX VX2 gel documentation system (Peqlab).

### Pulse Labeling

Yeast strains were grown either in full or selective media at 30 °C, 220 rpm to an OD_600_ of 0.5–1. Cells were harvested at 900 x g, RT for 5 min, washed twice with Labeling Medium (0.67% YNB without amino acids and ammonium sulphate/5% glucose, supplements as required by the strain’s auxotrophies), and resuspended in Labeling Medium to an OD_600_ of 6. Aliquots of 1.5 OD_600_ were transferred to clean 2 ml microfuge tubes. The samples were pre-incubated at the respective temperature, 800 rpm for 10 min to use up intracellular methionine and cysteine. Cells were then pulsed with 2.20 MBq per sample with Express Protein Labeling Mix (Perkin Elmer) and incubated for 2.5, 5, or 15 min (depending on the substrate) at 800 rpm, at the respective temperature. Cells were immediately transferred to ice and killed by adding 750 μl of cold Tris-Azide Buffer (20 mM Tris-HCl, pH 7.5/20 mM sodium azide). Cells were harvested for 1 min at full speed in a 5424-R Eppendorf microfuge at 4 °C, the pellets were resuspended in 1 ml of Resuspension Buffer (100 mM Tris-HCl, pH 9.4/10 mM DTT/20 mM ammonium sulphate) and incubated for 10 min at RT. The samples were centrifuged as before and resuspended in 150 μl of Lysis Buffer (20 mM Tris-HCl, pH 7.5/2% SDS /1 mM PMSF/1 mM DTT). Glass beads (150 μl, acid washed, 1 mm, Sigma) were added and the cells were disrupted in a MiniBeadbeater-24 (Bio Spec Products Inc.) for 2 x 1 min with 1 min pause in between cycles at RT. Samples were denatured at 85 °C for 5 min (10 min at 65 °C for membrane proteins). Beads were washed 3 times with 250 μl of IP Buffer without SDS (15 mM TrisHCl, pH 7.5/150 mM NaCl/1% Triton X-100/2 mM sodium azide), and the combined supernatants from each sample were collected and submitted to immunoprecipitation.

### Immunoprecipitation

Samples were precleared by adding 60 μl of 20% Protein A Sepharose™ CL-4B (GE Healthcare) in IP Buffer (15 mM Tris-HCl, pH 7.5/150 mM NaCl/1% Triton X-100/2 mM sodium azide/0.1% SDS) incubating for 30 min under rotation at RT. Samples were centrifuged for 1 min at full speed at RT and each supernatant was transferred to a clean microfuge tube containing 60 μl of 20% Protein A Sepharose™ CL-4B as well as the appropriate antibody. The samples were then incubated overnight at 4 °C under rotation. Samples were centrifuged for 10 sec at full speed, RT, washed with 1 ml of IP Buffer with SDS and 1 ml of Urea buffer (2 M Urea/200 mM NaCl /1 % Triton X-100/100 mM Tris-HCl, pH 7.5/2 mM sodium azide) 2 times each, and washed once with 1 ml of ConA buffer (500 mM NaCl/1 % Triton X-100/20 mM Tris-HCl, pH 7.5/2 mM sodium azide) and 1 ml of Tris-NaCl Wash (50 mM NaCl/10 mM Tris-HCl, pH 7.5/2 mM sodium azide). Samples were centrifuged as before and the supernatants discarded. SDS-PAGE Protein Sample Buffer (25 μl of 2x, 125 mM Tris-HCl, pH 6.8/4% SDS /10% β-Mercaptoethanol/0.002% bromophenol Blue/20% glycerol) was added and the samples incubated at 95 °C for 5 min (10 min at 65 °C for membrane proteins). Samples were loaded onto a 10% or 7.5% Bis-Tris gel (Invitrogen) and, following the electrophoresis, gels were fixed (10% acetic acid /40% methanol) for 30 min under shaking. After washing with deionized water, gels were dried at 80 °C for 1 h in a gel dryer (Model 583, Bio-Rad), exposed to phosphorimager plates and signal acquired in Typhoon Trio™ Variable Mode Imager (GE Healthcare). Signals were analysed and quantified using the ImageQuant™ TL software (GE Healthcare).

### Creation of libraries for screening and high content screen

To create the libraries, query strains that were generated as follows: KRY1160 and KRY1169 (KRY1160 transformed with pRS415 *sbh1S3A/T5A*) and crossed against the mini-Secretome-GFP library (382 proteins; *Geva et al., 2017*) using automated mating approaches (*Cohen and Schuldiner, 2011*; *Tong et al., 2001*). Then final collections were imaged by high-throughput fluorescence microscopy and correspondent image analysis was performed as described extensively (*Breker et al., 2014*).

## ACKNOWLEDGMENTS

We thank Nathan Ribot, Paula Hahn and Antinea Barbarit (all former students of the Römisch lab) for their contributions to this work as follows: Nathan Ribot for the construction of KRY1169. Paula Hahn for the screen through loss of function mutants in proline-directed kinases in yeast using the Sbh1(Pi) antibody vs. the Sbh1(10-23) antibody. Antinea Barbarit for the experiment shown in Fig. 1D. We thank Aline Leguede for expert technical assistance, Mark Lommel for help with strain construction and critically reading the manuscript, and Lihi Gal (Schuldiner lab) for help in high content screening. Work in the Schuldiner lab for this manuscript is part of a project that has received funding from the European Research Council (ERC) under the European Union’s Horizon 2020 research and innovation program (EU-H2020-ERC-CoG; grant name OnTarget, grant number 864068 to M.S.). M.S is an incumbent of the Dr. Gilbert Omenn and Martha Darling Professorial Chair in Molecular Genetics. The robotic system of the Schuldiner lab was purchased through the kind support of the Blythe Brenden-Mann Foundation. We thank Tom Stevens for the anti-DPAPB antibody; Howard Riezman for the anti-Gas1 antibody; and Charles Barlowe for the anti-Gls1 antiserum and the full-length GST-GLS1 expression construct.

## AUTHOR CONTRIBUTIONS

**Table EV1.**
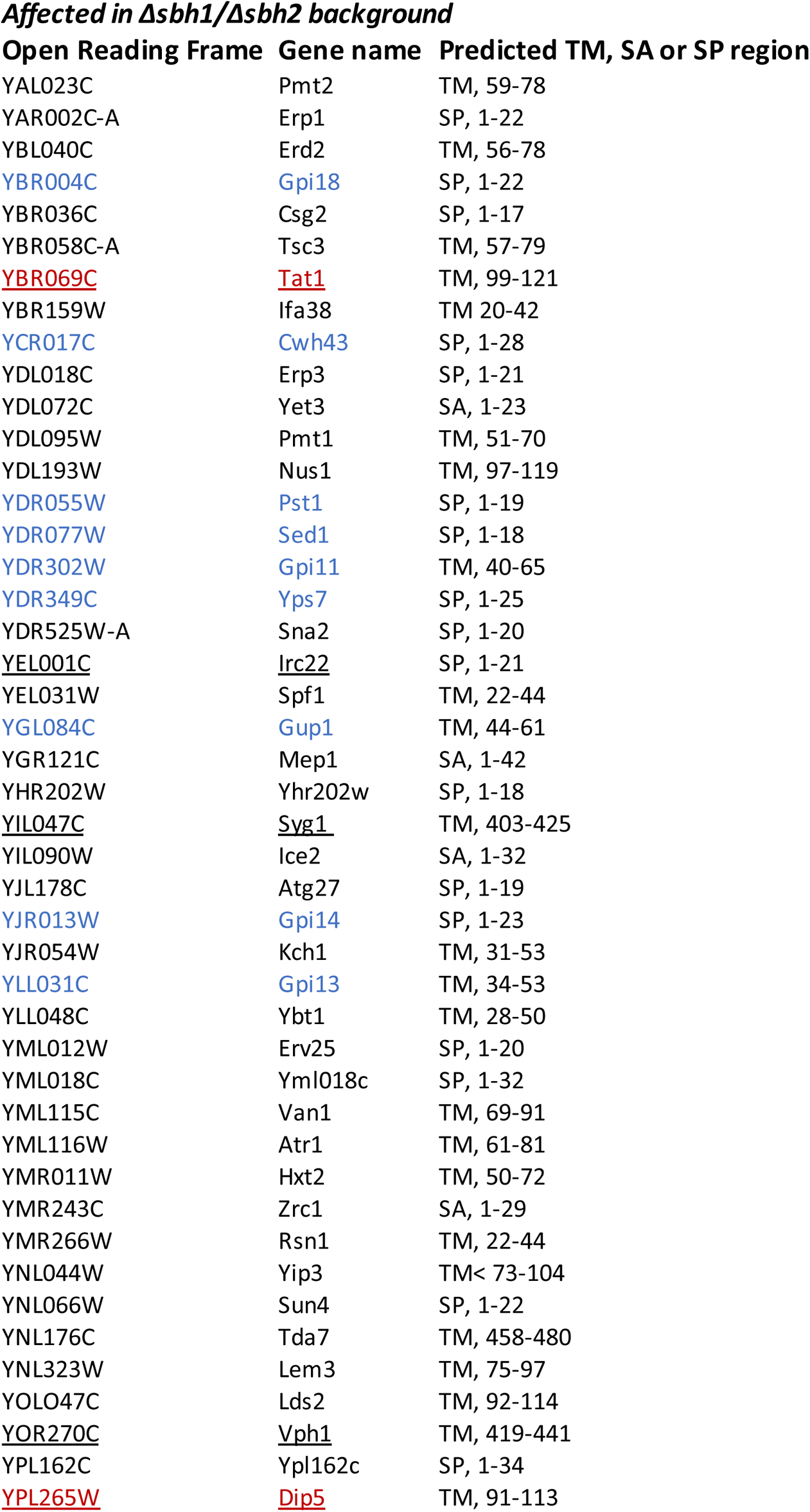
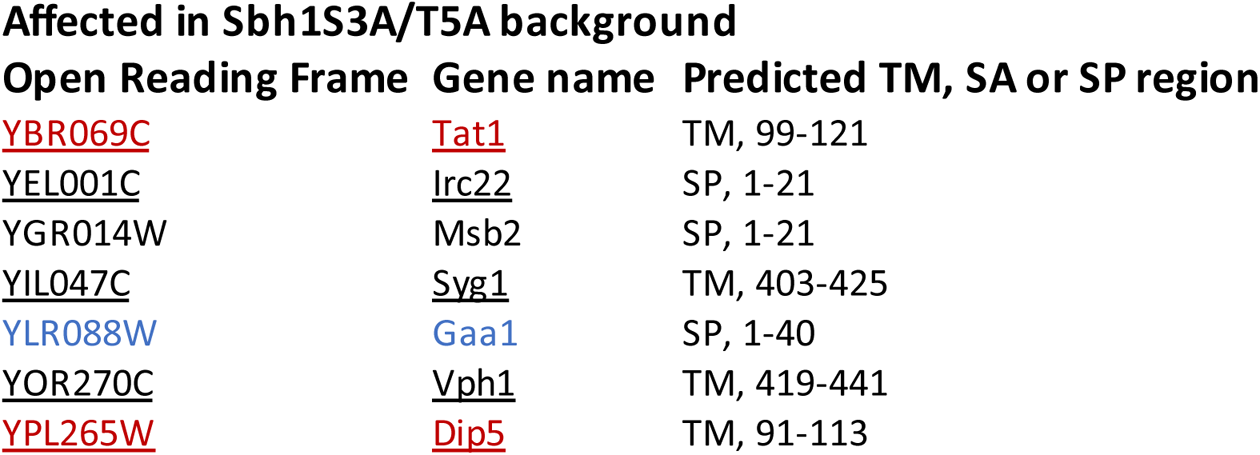
The Sbh1-dependent and Sbh1 phosphorylation-dependent ER translocation substrates from the high content screen. Blue: proteins involved in cell wall biosynthesis, red: amino acid transporters, underlined: overlapping proteins between the two screens. If the substrate is predicted to have a signal peptide (SP), a signal anchor (SA) or transmembrane targeting sequence (TM) and the predicted position of it, is also indicated.

**Table EV2.**
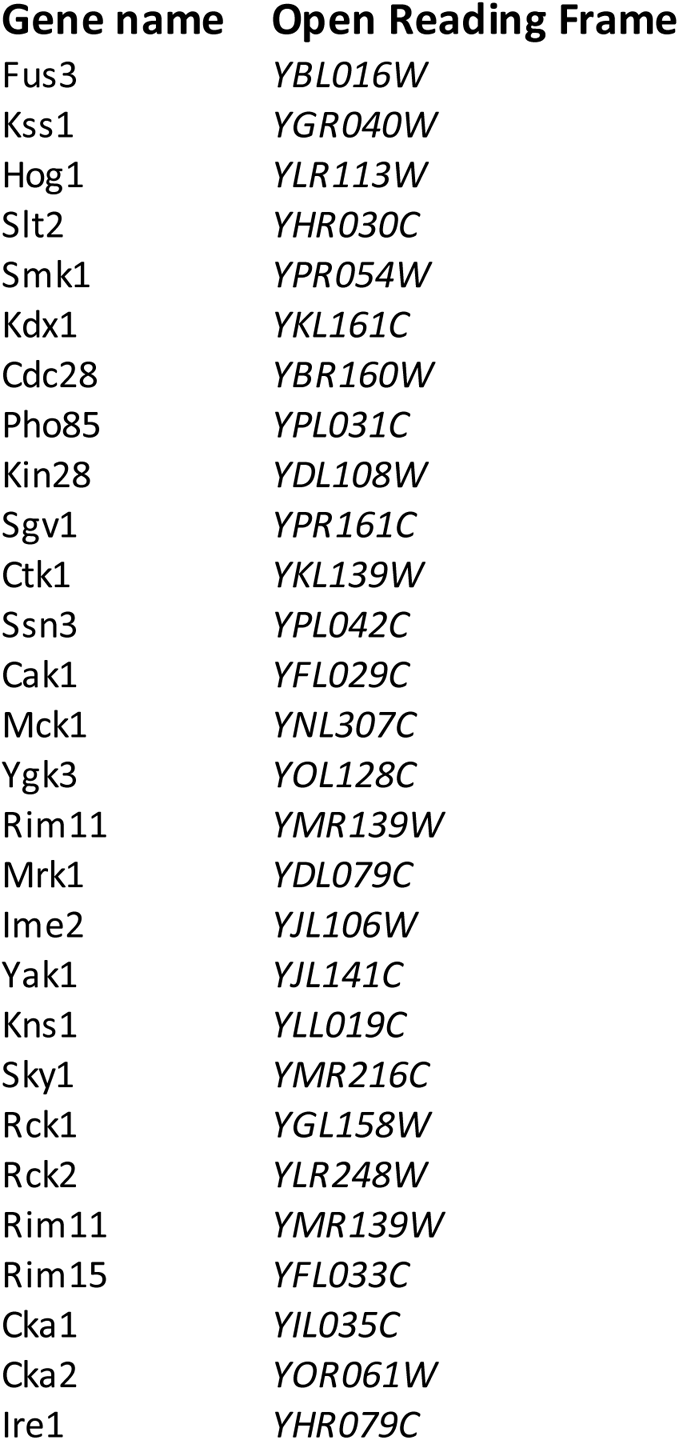
**Proline-directed Kinases:** The 27 different proline-directed kinases that are potentially responsible for the S3/T5-Sbh1 phosphorylation, and were tested in different screens. Ire1 is only known to phosphorylate itself so far, but since it associates with the Sec61 translocon we included it in the screen.

## EXPANDED VIEW FIGURE LEGENDS

**Figure EV1.**
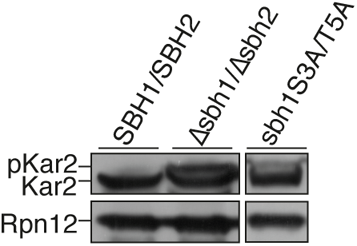
Cellular protein was extracted from wildtype, *Δsbh1Δsbh2* and *sbh1S3A/T5A* cells and Kar2 was detected by Western blotting with specific antibodies. Rpn12 was used as loading control.

**Figure EV2.**
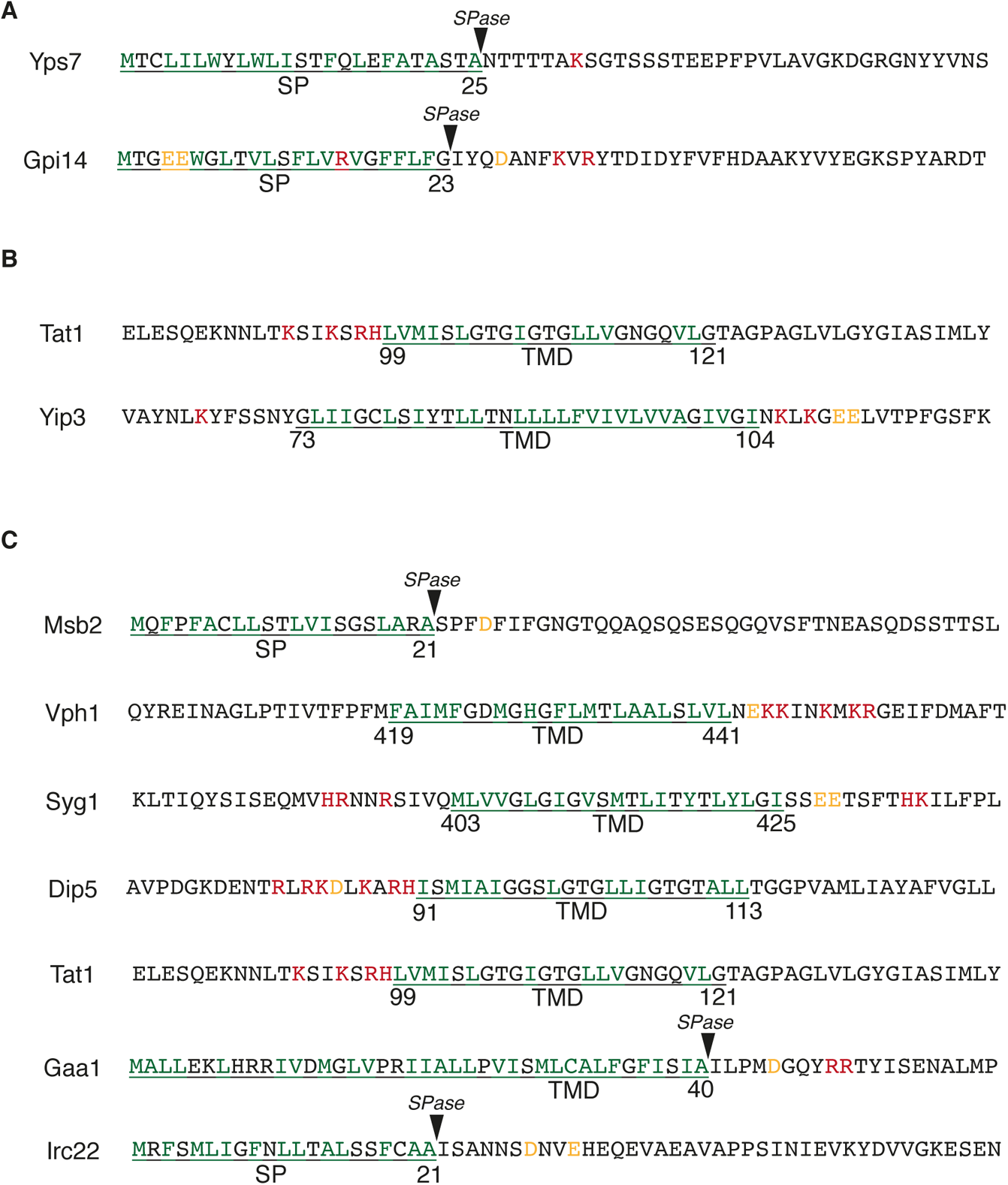
**(A)** Examples of signal peptide (SP) or first transmembrane domain (TMD) and the neighbouring sequences from the Sbh1-dependent ER translocation substrates from the high content screen with either no charge bias or an inverse charge bias. Underlined: SP or first TMD form the substrates, green: Hydrophobic residues from the predicted SP or TMD, red: Positive charged residues from the SP or TMD 10 residues neighbouring sequences, yellow: negatively charged residues from the SP or TMD 10 residues neighbouring sequences. Signal peptidase cleavage site (SPase) is also shown. **(B)** Examples of the first TMD and the neighbouring sequences from the Sbh1-dependent ER translocation substrates from the high content screen with unusually long or short targeting sequences or a high number of glycine residues. Underlined: first TMD form the substrates, green: Hydrophobic residues from the predicted TMD, red: Positive charged residues from the TMD 10 residues neighbouring sequences, yellow: negatively charged residues from the TMD 10 residues neighbouring sequences. **(C)** The SP or first TMD and the neighbouring sequences from the Sbh1 phosphorylation-dependent ER translocation substrates from the automated microscopic screen. Underlined: SP or first TMD form the substrates, green: Hydrophobic residues from the predicted SP or TMD, red: Positive charged residues from the SP or TMD 10 residues neighbouring sequences, yellow: negatively charged residues from the SP or TMD 10 residues neighbouring sequences. Signal peptidase cleavage site (SPase) is also shown.

**Figure EV3.**
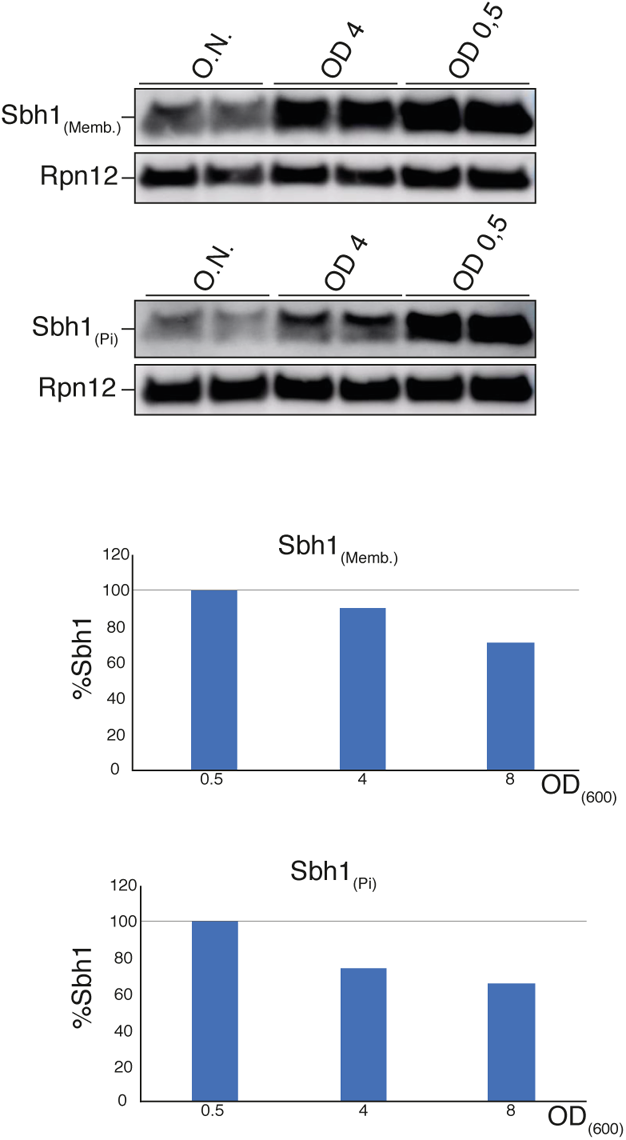
Cellular protein was extracted from wildtype yeast cells at different growth stages and Sbh1 and phosphorylated (Pi)-Sbh1 were detected by Western blotting with antibodies against the membrane-proximal conserved region without phosphorylation sites (amino acids 39-54) (upper) or the phosphorylated N-terminus (lower). Rpn12 was used as loading control.

**Figure EV4.**
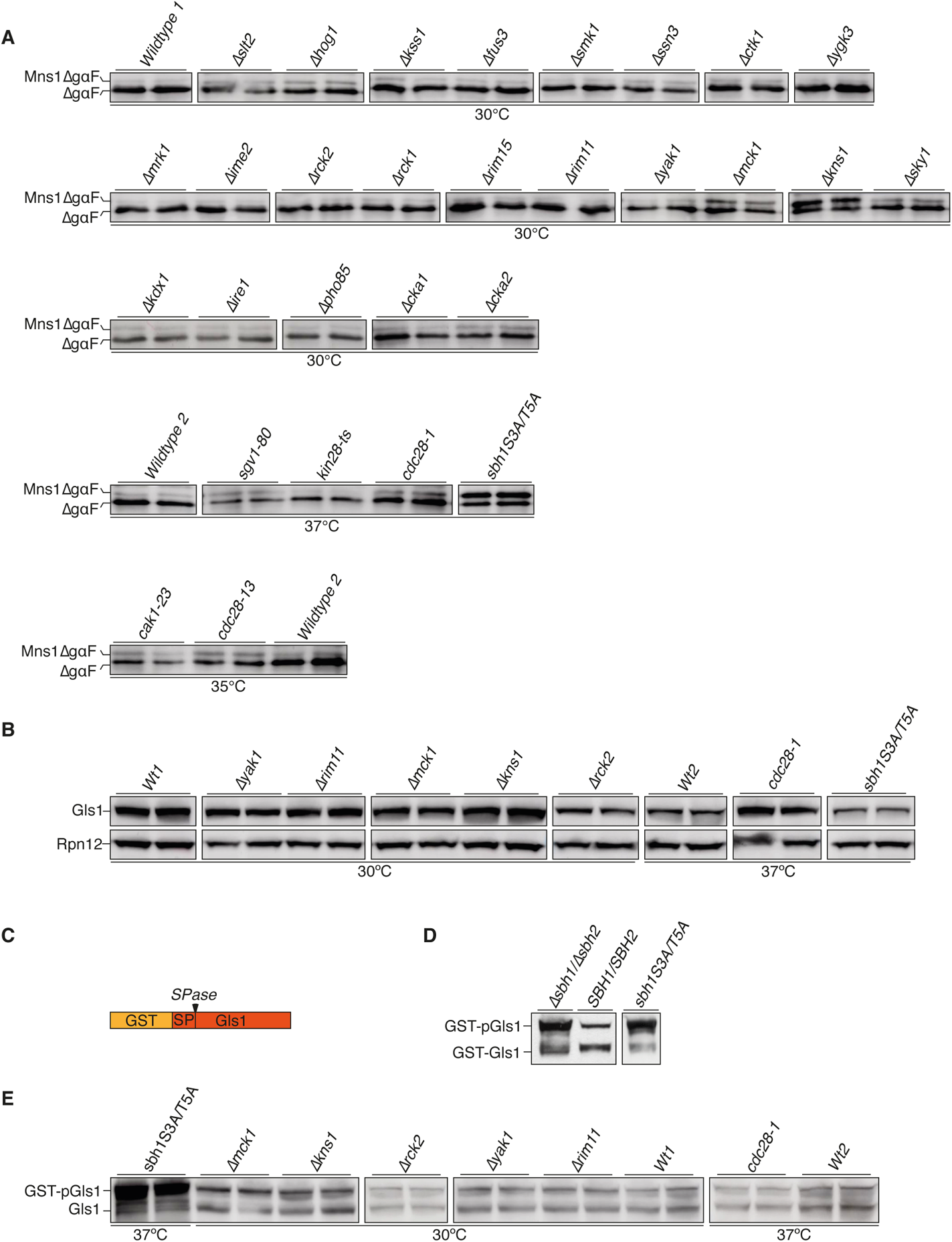
**(A, B)** Cellular protein was extracted from wildtype and the indicated mutant cells and Mns1ΔgpαF, Gls1, Rpn12 (loading control) were detected by Western blotting with specific antisera. **(C)** Schematic representation of GST (orange) fused to pGls1 (red). **(D, E)** Cellular protein was extracted from wildtype and the indicated mutants and GST-Gls1 was detected by Western blotting with Gls1 specific antibodies. Rpn12 was used as loading control. Cytosolic precursor (GST-pGls1) and ER form (Gls1) are indicated.

**Figure EV5.**
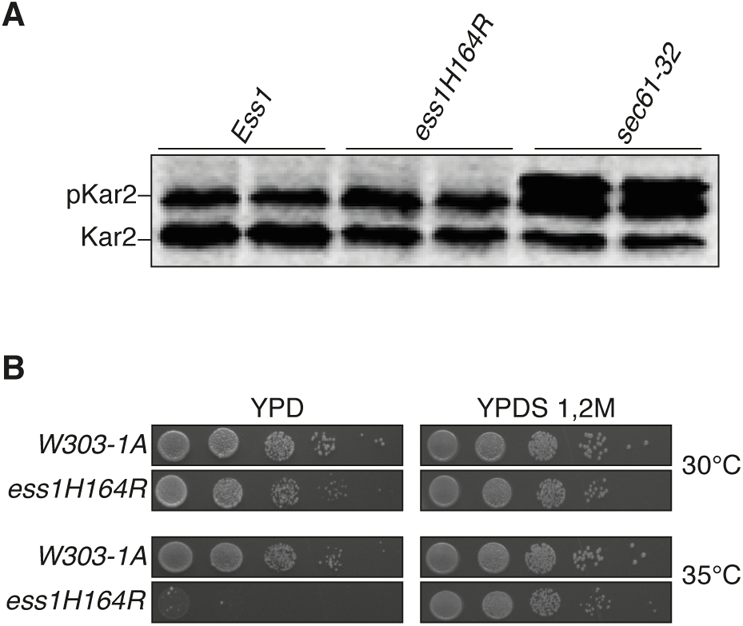
Wildtype and the indicated mutant strains were grown in serial dilutions on YPD and YPD+1.2 M sorbitol (YPDS) for 3 days at 30 °C and 35 °C.

